# The metallome of *Methanosarcina barkeri* during electron uptake from a cathode

**DOI:** 10.64898/2026.02.14.705942

**Authors:** Abdalluh Jabaley, Thor Frickman, Pia Bomholt Jensen, Mads Lykke Justesen, Daniel M. Chevrier, Damien Faivre, Thomas Boesen, Amelia-Elena Rotaru

## Abstract

Electromethanogenesis, the cathode-dependent reduction of CO_2_ to CH_4_ by methanogens, offers a sustainable route to methane fuel. *Methanosarcina barkeri* lacks surface-exposed multiheme cytochromes for extracellular electron transfer (EET). Instead, we recently showed that surface-bound G-quadruplex ribonucleic acids (G_4_-RNA) are required for EET, yet how electrons traverse this extracellular matrix remains unresolved. Here, we quantified metal accumulation during cathodic growth by inductively coupled plasma mass spectrometry in cells grown on cathodes poised at –430 mV versus the standard hydrogen electrode, using acetate-grown cells, open-circuit controls and abiotic cathodes for comparison. Cathode-grown *M. barkeri* showed CH_4_ buildup attributable to cathodic electrons (2.1 ± 0.8% CH_4_), whereas open-circuit controls showed negligible increase (0.26 ± 0.15% CH_4_). Cathode-bound cells exhibited ∼55-fold enrichment in Co, Ni and Mo, and 5- to 21-fold enrichment in Cu, Zn, and Fe relative to acetate-grown cells; neither acetate-grown cells nor abiotic cathodes accumulated metals. To resolve where metals reside, we mapped the elemental distribution in acetate-grown cells by scanning transmission electron microscopy-energy dispersive X-ray spectroscopy and high-resolution nano X-ray fluorescence. Fe co-localized with phosphorus in intracellular storage bodies, whereas Co and Zn localized within the extracellular capsule. Together, these data indicate selective metal sequestration during electromethanogenesis and raise the possibility that certain metals associate with G_4_-RNA and/or the methanochondroitin matrix to support charge transfer at the cell surface. This metalomic fingerprint provides a new proxy for dissecting archaeal EET strategies and may inform the design of more efficient bioelectrochemical systems.

## Introduction

Methanogens drive the final step of anaerobic organic matter degradation, and are the major biological drivers of global methane emissions as well as key players in anaerobic biogas digester technologies[1, 2]. Among methanogens, *Methanosarcina barkeri* stands out for its metabolic versatility: it can employ all known methanogenic pathways, including methylotrophic, acetoclastic and CO_2_ reduction with H_2_ [3] or with electrons directly provided by other cells or electrodes[4, 5]. The latter makes *M. barkeri* an ideal model organism for studying extracellular electron transfer (EET) and its applications in bioelectrochemical technologies such as electromethanogenesis, the sustainable conversion of CO_2_ and renewable electricity to methane fuel[2, 6].

Extracellular electron transfer has been primarily studied in model electrogens, such as *Geobacter* species, which rely on extracellular multiheme c-type cytochromes (MHCs) either embedded in extracellular polymeric substances (EPS), bound to conductive pili or forming own nanowire-structures to reach faraway extracellular electron acceptors[7–13]. Unlike these bacteria or even closely related methanogens like *M. acetivorans*[14–16], *M. barkeri* lacks the canonical structures for EET[17]. Instead, *M. barkeri* grows in large multicellular aggregates encased by a unique extracellular polysacharide, methanochondroitin, composed of repeating trimers of N-acetylglucosamine and glucuronic acid in the configuration [→4)-β-D-GlcUA(1→3)-D-GalNAc-(1→ 4)-D-GalNAc(1 →]n [18].

The ability of EPS-matrices to conduct electrical current has been linked to their capacity to bind molecules that can support charge transfer (e.g., metals, flavins, hemins, cytochromes)[19, 20]. In several bacteria, extracellular matrices are composed not only of EPS but also of extracellular nucleic acids (e.g., eNAs), some of which form G-quadruplex (G_4_) structures stabilized within the EPS[21]. In biological systems, G_4_-NAs are capable of conducting electrical charge when coordinated with metals or redox-active cofactors (e.g., hemin, phenazines)[22, 23]. For example, *Staphylococcus epidermidis* forms electroactive biofilms in the presence of alkali-metal-stabilized G_4_-DNA structures and hemin[23]. In *M. barkeri*, we recently demonstrated that self-released extracellular G_4_-RNAs are required for EET: their depletion abolishes cathodic electron uptake, whereas supplementation enhances electromethanogenesis[24]. These findings raise the possibility that the extracellular matrix of *M. barkeri* provides a structural scaffold for organizing charge-transport constituents on the cell surface.

Metal dependence is universal among methanogens, as they share many metalloenzymes essential for methanogenesis, including Ni in methyl-CoM reductase, Co in corrinoid methyltransferases, Mo in formyl-methanofuran dehydrogenase[25], and multiple Fe–S cluster-containing enzymes such as heterodisulfide-reductase [26]. However, *M. barkeri* is metabolically versatile, capable of employing multiple methanogenic pathways, which may impose increased demands on metal acquisition and allocation. In addition, unlike other methanogens, *M. barkeri* poseses a distinctive cell surface architecture and can receive extracellular electrons, potentially creating unique metal requirements at the cell-electrode interface. Although, trace metal supplementation enhances methane yields in mixed-community digesters [27], metal homeostasis in methanogen monocultures under controlled electrochemical conditions remains unexplored. In hypothesized that, in the absence of multiheme cytochromes, *M. barkeri* leverages it’s cell surface architecture to sequester metals and confactors not only for intracellular functions but also possibly to influence extracellular electron transfer.

In this study, we directly compare the metallome of *M. barkeri* during growth on cathodes poised at –430 mV vs the standard hydrogen electrode (SHE) with acetate-grown cultures and abiotic controls. We combine bulk metallome determination via inductively coupled plasma mass spectrometry (ICP-MS) with nanoscale elemental mapping by scanning transmission electron microscopy-energy dispersive X-ray spectroscopy (STEM–EDS) and synchrotron nano X-ray fluorescence (nano-XRF) to quantify and locate key metals (Fe, Ni, Co, Mo, Zn, Cu). Our objective was to determine whether cathodic growth triggers selective metal enrichment in cathode-bound cells. We also mapped baseline metal localization in acetate-grown cells. Our results show selective metal enrichment during electromethanogenesis and establish where specific metals typically reside in *M. barkeri* cells.

## Methodology

### Cultures, media and growth conditions

*Methanosarcina barkeri* MS (DSM800) was obtained from the German Collection of Microorganisms and Cell Cultures (DSMZ, Germany). Cultures were grown in modified DSM-120 medium lacking resazurin, sodium sulphide, yeast extract and tryptone[28]. All media and cultures were prepared and maintained under strictly anaerobic conditions with an N_2_:CO_2_ (80:20, v/v) headspace and incubated at 37°C. Routine transfers used 10% (v/v) mid-exponential phase culture as inoculum.

Three experimental conditions were used: (i) bioelectrochemical reactors with cells (biotic), (ii) bioelectrochemical reactors without cells (abiotic controls) and (iii) acetate-grown cells (acetoclastic) as additional controls.

Acetoclastic cultures were prepared anaerobically as previously described[29] in six 100 mL replicates, with 50 mM acetate as substrate. Bioelectrochemical reactors were operated either under poised conditions (-430 mV vs SHE) or under open-circuit conditions. In open-circuit conditions, electrodes were present but not connected (no current flow). Biotic reactors were run in three poised biological replicates (n=3) and four open circuit biological replicates (n=4). Abiotic controls consisted of six poised replicates (n=6) and three open circuit replicates (n=3).

Reactors were gas-tight H-cell systems (Adams and Chittenden Scientific Glass, USA) composed of two conjoined chambers separated by a Nafion-N117 proton-exchange membrane (Ion Power, Germany), activated in deionized water at 60 °C for 50 min. A three-electrode configuration was used, with carbon felt (1.27 x 2.5 x 2.5 cm, Thermo Scientific, USA) as working and counter electrodes connected via titanium wires. A leak-free Ag/AgCl reference electrode (3.4 M KCl, CMA Microdialysis, Sweden) was positioned ∼1 cm from the working electrode in the cathodic chamber. Electrode wires were then connected to a MultiEmstat potentiostat (PalmSens, The Netherlands). Electron uptake was monitored using chronoamperometry and methane production was confirmed by gas chromatography (GC) of headspace samples. All reactors were inoculated with 20% (v/v) culture (OD600nm ∼0.12). The inoculum was pre-grown on 10 mM methanol and 15 mM acetate and reached the required optical density after 72 h. Previous tests confirmed that acetate carryover did not exceed 2.5 mM, which was insufficient to sustain detectable acetoclastic methanogenesis from acetate. Reactors were incubated at 37 °C for 14 days in 37 °C with continuous stirring (100 rpm).

### Protein extraction and quantification

For acetoclastic incubations, proteins were determined alongside methane, acetate, optical density, and metal content at each sampling point (days 0, 3, 6, 10, 13, 18, 24).

Bioelectrochemical reactors were disconnected and disassembled after 14 days. The carbon cathodes were rinsed with fresh DSM120 media to remove loosely attached cells, and then cut into equal pieces (1.27 x 0.5 x 0.5 cm) to obtain technical replicates from each cathode for statistical analysis. Three pieces per cathode were transferred to separate Eppendorf tubes containing 1 mL of a CHAPS solution (2 % w/v in phosphate buffer saline, pH = 7.4), then boiled for 5 min to release proteins from the electrode surface. Extracts were centrifuged at 16,000 g for 15 min at 4 °C, and the supernatants were stored at –80 °C until all were collected for protein determination. Protein concentration was determined using the bicinchoninic acid (BCA) assay according to the manufacturers instructions (Novagen, USA) and measured on a Multiscan Go (Thermo Scientific) spectrophotometer at 562 nm.

### Analytical measurements

Acetate samples from acetoclastic incubations were retrieved aseptically and anaerobically on days 0, 3, 6, 10, 13, 18, and 24. Acetate concentrations were quantified using ion chromatography as previously described[30]. Methane was sampled aseptically and anaerobically and quantified by gas chromatography as previously described[30]. In brief, 100µL of headspace gas from each incubation was injected into a Trace1300 (Thermo Fisher Scientific, USA) gas chromatograph equipped with a TG-BOND Sieve 5A column (Thermo Fisher Scientific, USA) and a flame ionization detector (FID). The carrier gas was N_2_. The injector, oven, and detector temperatures were 190, 100, and 200 °C, respectively.

### Sampling and metal determination by ICP-MS

For metal determination from acetate-grown cells, 1 mL culture was harvested at each sampling point by centrifugation (7,000 x g, 10 min). The supernatant was removed, and the pellets were suspended in 1 mL of 2% (v/v) HNO_3_. For metal determination on electrodes, after 14 days, cathodes (biotic and abiotic) were washed with fresh DSM120 medium and cut into equal pieces (1.27 x 0.5 x 0.5 cm). Three pieces of each cathode were used for technical replication; each piece was placed in 1 mL of 2% (v/v) HNO_3_. The samples then went through three cycles of freeze-thawing (liquid nitrogen and 60 °C) to lyse the cells. Finally, to remove any carbon cloth particles, the liquid phase containing extracted metals was filtered (0.45 µm). The metal samples were then diluted 20-fold prior to total metal concentrations being determined by inductively coupled plasma mass spectrometry (Agilent 7900 ICP-MS) as previously described[31].

### Electron microscopy

Mid-exponential, acetate-grown *M. barkeri* cells (5 mL) were harvested by centrifugation (7,000 x g, 10 min). The supernatant was removed, and pellets were fixed in 250 µL fixative containing 2.5% (v/v) formaldehyde and 2.5% (v/v) glutaraldehyde in 0.1 M sodium cacodylate buffer (pH 7.4) (Electron Microscopy Sciences, USA). Samples were stored in fixative at 4 °C until processing. Scanning electron microscopy (SEM) imaging was performed using a Zeiss Sigma 300 VP. Fixed *M. barkeri* cells were deposited directly onto an Al stub and coated with ∼10 nm carbon prior to imaging. Imaging was conducted using the InLens detector under high vacuum (1 × 10^−6^ bar) at 1 kV. For downstream STEM-EDS, fixed cells were embedded in molten agar, and after solidification, embedded in Epon, as previously described[32]. In brief, the agar blocks were placed in silicone molds, then Epon was poured and cured at 60 °C for 24 h. The Epon-blocks were sectioned into 100 nm thin sections using an Ultramicrotome EM UC7 (Leica). The thin sections were mounted carbon-coated copper grids (collodion support film) that were glow discharged prior to use.Thin sections of fixed cells were negatively stained [33] and examined by transmission electron microscopy (TEM) (Tecnai G2 Spirit), while no negative stain was applied for STEM using a Talos F200X (S)TEM (Thermo Fisher Scientific). Scanning transmission electron microscopy with energy-dispersive X-ray spectroscopy (STEM-EDS) was performed using a FEI Talos FX200i STEM (Thermo Fisher Scientific). Spectrum images were aquired over 120-240 min using a beam current of 7.48 nA, a beam voltage of 200 keV, and a dwell time of 20 µs. Velox software (Thermo Fisher Scientific) was used to process spectrum images and generate elemental maps.

### Nanobeam-scanning X-ray fluorescence microscopy

Measurements of a fraction of interstitial cell material (possibly consisting of methanochondoitin and other extracellular substances) were conducted at the I14 nanoprobe of Diamond Light Source (Oxfordshire, UK). Samples were measured in dried condition on copper electron microscopy grids. All measurements were conducted under ambient pressure and temperature using an incident photon energy of 11.5 keV. The focused X-ray beam was ∼50 nm (FWHM) in diameter. X-ray fluorescence (XRF) was collected from the front of the sample using a four-element silicon drift detector (RaySpec, UK)[34]. A raster scanning step size of 50 nm was used with an exposure time of 50 ms per pixel to collect high-resolution XRF maps. A photon-counting Merlin detector (Quantum Detectors, UK) was used in transmission configuration to collect X-ray scattering for generating phase gradient images. Detailed information on how the transmitted signal was transformed, including masking of the beam, background intensity adjustment, and phase integration, is described by Quinn et al[34] PyMCA 5.6.7 software was used to energy calibrate XRF spectra, background subtraction, fit XRF sum-spectra and export XRF maps for individual emission lines (e.g., Ca Kα). ImageJ was used to further render XRF maps.

## Results and discussion

### Electromethanogenesis is associated with selective metal enrichment on cathodic M. barkeri

When presented with a carbon-felt cathode poised at -430 mV vs SHE, *M. barkeri* drew a sustained cathodic current that was absent from cell-free abiotic cathodes (Fig. 1a), consistent with cellular electron uptake as reported previously for this methanogen[4, 5]. After 15 days, *M. barkeri* in poised reactors accumulated 2.1 ± 0.8 % CH_4_ (n=3) compared to 0.26 ± 0.15% in open-circuit controls (n=4, p=0.007) (Fig. 1b). If methane production was driven by hydrogenotrophic methanogenesis, it would require substantial H_2_ availability (4 mol H_2_ per mol CH_4_). However, in abiotic reactors operated at -430 mV over a 15-day period, H_2_ remained near the detection limit (0 to 0.07%; mean 0.03 ± 0.03% H_2_), which could support at most 0 to 0.015% CH_4_, assuming complete stoichiometric conversion, and without considering the relatively high H_2_ thresholds reported for *Methanosarcina* spp. [35, 36]. The lack of electrochemically generated H_2_ is consistent with previous measurements, and with linear sweep voltammetry, which in this setup places the onset of abiotic H_2_ evolution at substantially more negative potentials (≤ -700 mV vs SHE)[5]. Together these observations indicate that abiotic H_2_ evolution is negligible at -430 mV and cannot account for the bulk methane production in the poised reactors.

**Figure 1.**
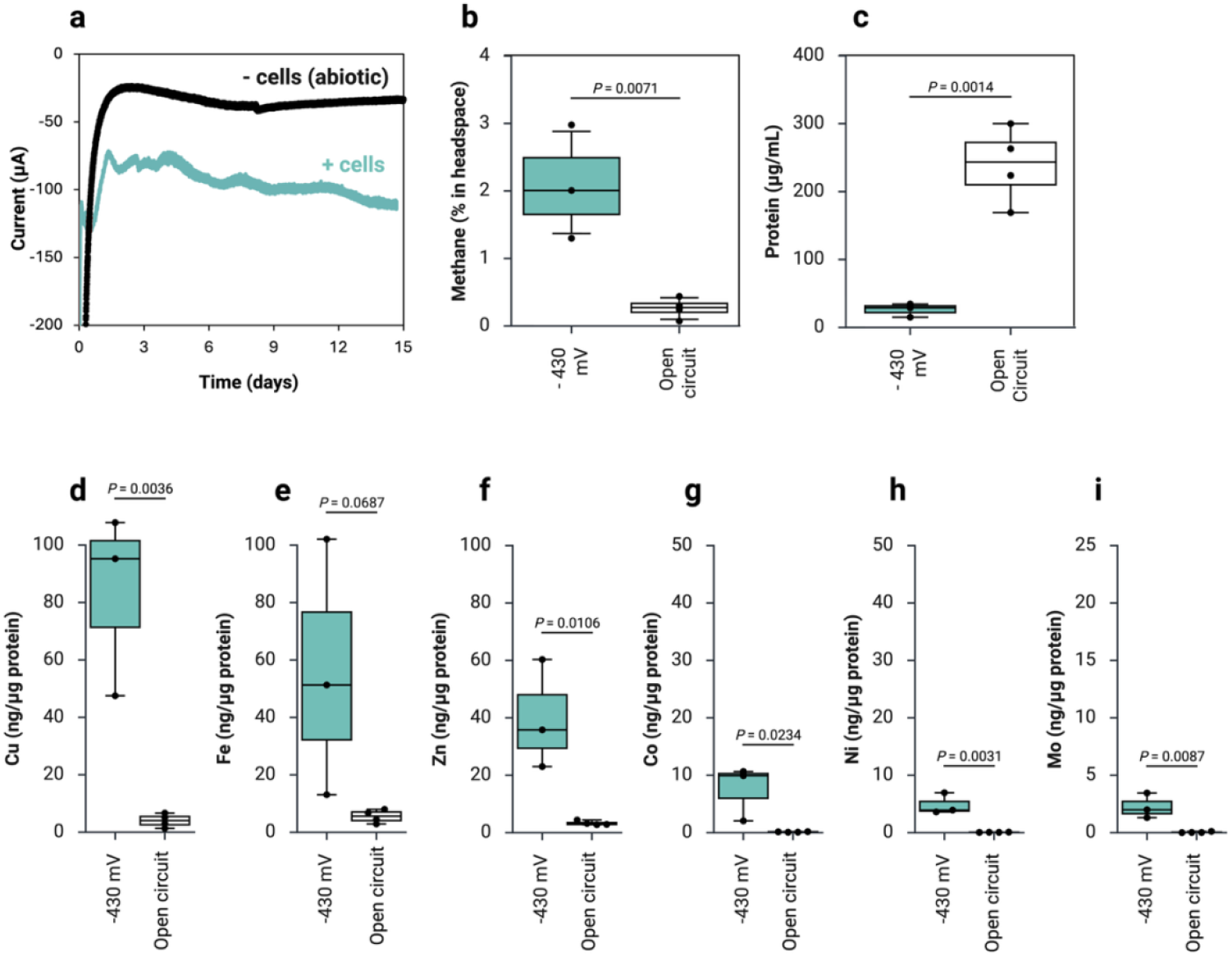
Electron uptake and methane production by *Methanosarcina barkeri* on cathodes poised at -430 mV versus open circuit (disconnected) controls, with associated protein and metal content. (a) Chronoamperometry of cathode-fed *M. barkeri* (blue-trace) and abiotic controls (black trace). (b) Final methane yields after 15 days under poised and open circuit conditions (n=3 and n=4 respectively). (c) Total protein recovered from carbon cloth electrodes on day 15 under poised and OC condtions (n=3 and n=4 respectively). (d-i) Fe, Cu, Zn, Co, Ni, and Mo content of biocathodes normalized to protein content (ng metal per µg protein). Box plots show the median of minimum three biological replicates (horizontal line), interquartile range (box), and full range (whiskers); individual biological replicates are shown as open circles and each of the biological replicates is the median value of three independent technical replicates. Statistical significance between conditions was assessed using an unpaired t-test. Figure compiled in Biorender.

To determine whether metals accumulate in *M. barkeri* during electromethanogenesis, we quantified the metal content in cathode-associated biomass from electrodes poised at -430 mV vs SHE and compared it to biomass recovered from electrodes under open circuit (OC) condition. Cells on poised cathodes accumulated substantially more metal per unit protein than cells on OC-control electrodes. Notably, total protein recovered from poised electrodes was lower than from OC-controls (Fig. 1c) consistent with reduced attachment and/or enhanced biofilm dispersion in microbial electrosynthesis, at negative electrode potentials[37]. Despite lower cathode-associated protein, *M. barkeri* in poised reactors produced methane and remained electroactive through the incubation period (Fig. 1a-b), indicating that reduced attached biomass does not reflect loss of metabolic activity.

On poised biocathodes the three most abundant metals were Cu, Fe and Zn. Among these, Cu was the most abundant (83.5 ± 31.8 ng Cu µg^-1^ protein; n=3), and was enriched ∼21-fold relative to OC-controls (4.0±2.3 ng Cu µg^-1^ protein; n=4; p=0.004) (Fig. 1d). Fe was the second most abundant and was enriched on average ∼10-fold relative to OC-controls (55.4 ± 44.6 vs 5.5 ± 2.3 ng Fe µg^-1^ protein), but this trend did not reach significance due to high between-replicate variability (p=0.069) (Fig. 1e). Zn was enriched ∼12-fold on poised biocathodes compared to OC-controls (39.7 ± 18.9 vs 3.3 ± 0.7 ng Zn µg^-1^ protein; p = 0.011).

Elevated Fe is consistent with the high Fe demand of methanogens, where energy metabolism relies heavily on Fe-S proteins[38, 39]. Besides, *M. barkeri* upregulates Fe-S proteins during direct interspecies electron transfer, suggesting that electron uptake metabolism depends on an Fe-rich electron transfer machinery[40].

Zn enrichment is also linked to energy metabolism in methanogens, having established roles in coenzyme M activation [41] and in the core methylotrophy machinery of *Methanosarcina*, including Zn-dependent corrinoid methyltransferases [42]. To assess whethes additional Zn enzymes could demand higher Zn uptake, we screened the *M. barkeri* genome and identified other predicted Zn proteins, including a γ-class carbonic anhydrase[43] and membrane-bound Zn-metalloproteases linked to stress response[44].

On the other hand, Cu is not a canonical cofactor of core methanogenesis and is tightly controlled due to its toxicity. It occurs in small copper proteins such as cupredoxins, where Cu centers mediate electron transfer[45]. Cu is highly toxic outside a narrow physiological range; for example, Cu inhibits *Methanospirillum hungatei* at just 50 µM [46], whereas *M. barkeri* tolerates Cu up to ∼1.5 mM without inhibition [47]. Because Cu is toxic, microorganisms regulate Cu influx and efflux at the cell surface through controlled transport and sequestration, including EPS binding and Cu-binding proteins that complex free Cu [48]. Accordingly, we found that *M. barkeri* encodes Cu/Zn transporters and a CopA-like Cu-translocating ATPase central to Cu homeostasis. In the biocathode context, Cu enrichment may therefore reflect altered Cu binding at the cell-electrode interface, either via increased abundance of Cu-binding proteins and/or sequestration of Cu within the methanochondroitin capsule and the G4 nucleic acids embedded in it [24]. Cu has been shown to influence G4-structures by stabilizing them, inducing specific folding topologies, and enabling conformational switching [49, 50]. Such Cu-induced structural changes could, in turn, modulate G4-mediated charge transfer and their role as EET conduits between the electrode and the cell [24].

Nevertheless, the most significant change in terms of enrichment on biocathodes and open circuit controls was observed for trace metals like Co, Ni and Mo. Co was the highest most abundant trace metal, enriched ∼63-fold on biocathodes than on OC-controls (7.5 ± 4.8 vs 0.12 ± 0.0, ng Ni µg^-1^ protein p=0.023). Ni was enriched ∼69-fold on biocathodes than on OC-controls (4.8 ± 1.8 vs 0.07 ± 0.0, ng Ni µg^-1^ protein; p = 0.003) and Mo ∼58-fold (2.3 ± 1.1 vs 0.04 ± 0.04 ng Mo µg^-1^ protein; p=0.009).

Co is central to methanogenesis because it forms the metal center of corrinoid (cobamide)-cofactors used by methyltransferases in methylotrophic pathways, and also in the core CO_2_-reduction pathway via the membrane bound methyl-H_4_MPT:coenzyme M methyltransferase (Mtr), which contains a cobamide prosthetic group and couples methyl-transfer to energy conservation[51, 52]. Corrinoid-dependent catalysis requires the cobalt to be in the super-reduced state Co(I). *M. barkeri* has an Fe-S protein (RamA) which catalyses the ATP-dependent activation of Co(II) to Co(I) [53]. We postulate that cathodic growth shifts cellular redox conditions, increasing the burden of maintaining corrinoids in the active state, and thereby increasing Co demand to sustain corrinoid-dependent methyl transfer during CO_2_-reducing electromethanogenesis.

Similar to Co, Ni enrichment on biocathodes is consistent with increased demand for cofactors central to methanogenic energy metabolism. In methanogens, Ni availability directly impacts methane formation through the key terminal enzyme - methyl coenzyme M reductase (Mcr), which requires the unique Ni-cofactor F_430_[36]. Consistent with this, *M. barkeri* has a relatively high Ni requirement, and CO_2_-reductive methanogenesis is inhibited under Ni limitation[36].

Mo is also central to energy metabolism in CO_2_ reducing methanogens. In *M. barkeri* the first step of CO_2_ reduction is catalyzed by a Mo/W-enzyme (molybdopterin) – formylmethanofuran dehydrogenase (Fmd/Fwd). The Mo cofactor in *M. barkeri* was identified as a molybdopterin guanine dinucleotide-type cofactor [54]. We also found that M. barkeri had the genetic capacity for Mo-cofactor assembly and Mo-uptake [55, 56]. Thus, higher Mo enrichment on *M. barkeri* on poised cathodes may reflect increased CO_2_-reduction pathway throughput during electrode driven methanogenesis.

Two controls were run alongside to distinguish electromethanogenesis-driven metal accumulation from abiotic deposition under electrochemical conditions and growth-related effects. First, we operated cell-free (abiotic) reactors with carbon felt cathodes poised at - 430 mV vs open circuit, to determine whether the applied potential promotes electroplating with metals available in the culture medium. No metals increased on abiotic cathodes under poised condition relative to OC-controls (Fig. 2), indicating that the potential alone does not drive metal deposition on the electrode. If anything, Co was lower on poised abiotic cathodes than on abiotic OC-controls (Fig. 2d) further arguing against any metal electroplating in our setup. Second, we quantified metal-quotas in acetate-grown *M. barkeri* cells to determine whether methanogenic growth over ∼2weeks inherently leads to metal enrichment per cell. In acetate-grown cultures, metal quotas did not increase with time, instead metal-per-unit-protein decreased as biomass accumulated (Fig. 3). Even at the highest metal quotas during acetate growth, metal levels remained orders of magnitude lower than in cathode-grown biomass (∼25—9500-fold lower, depending on the metal) (Fig. 1, Fig. 3). Together, these controls show that the elevated metal content observed on poised biocathodes cannot be explained by cathode electroplating or by biofilm growth-associated metal accumulation, but instead reflects a selective metal sequestration response linked specifically to cathode-driven methanogenesis.

**Figure 2.**
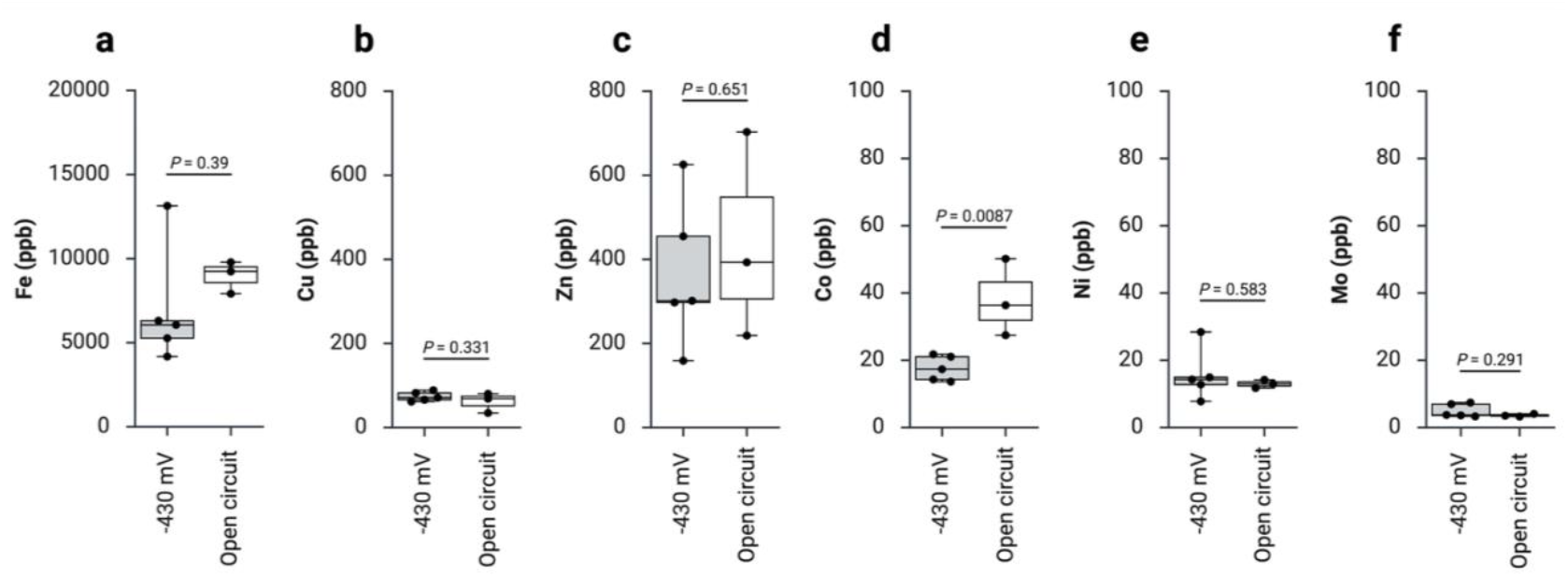
Metal levels on cell-free (abiotic) carbon-felt cathodes under poised (– 430 mV) and open-circuit (OC) conditions. Concentrations of Fe, Cu, Zn, Co, Ni, and Mo measured by ICP-MS and reported in parts per billion (ppb). All abiotic reactors were incubated for 15 days at 37 °C similar to biotic incubations (n=5 and n=3 for -430 mV versus OC). The median of quadruplicate technical replicates for each reactor replicate are shown as circles. On abiotic cathodes, unpaired t-tests reveal no significant difference between – 430 mV and OC for any metal other than Co, were less of the metal was detected under the poised condition. Figure compiled in Biorender.

**Figure 3.**
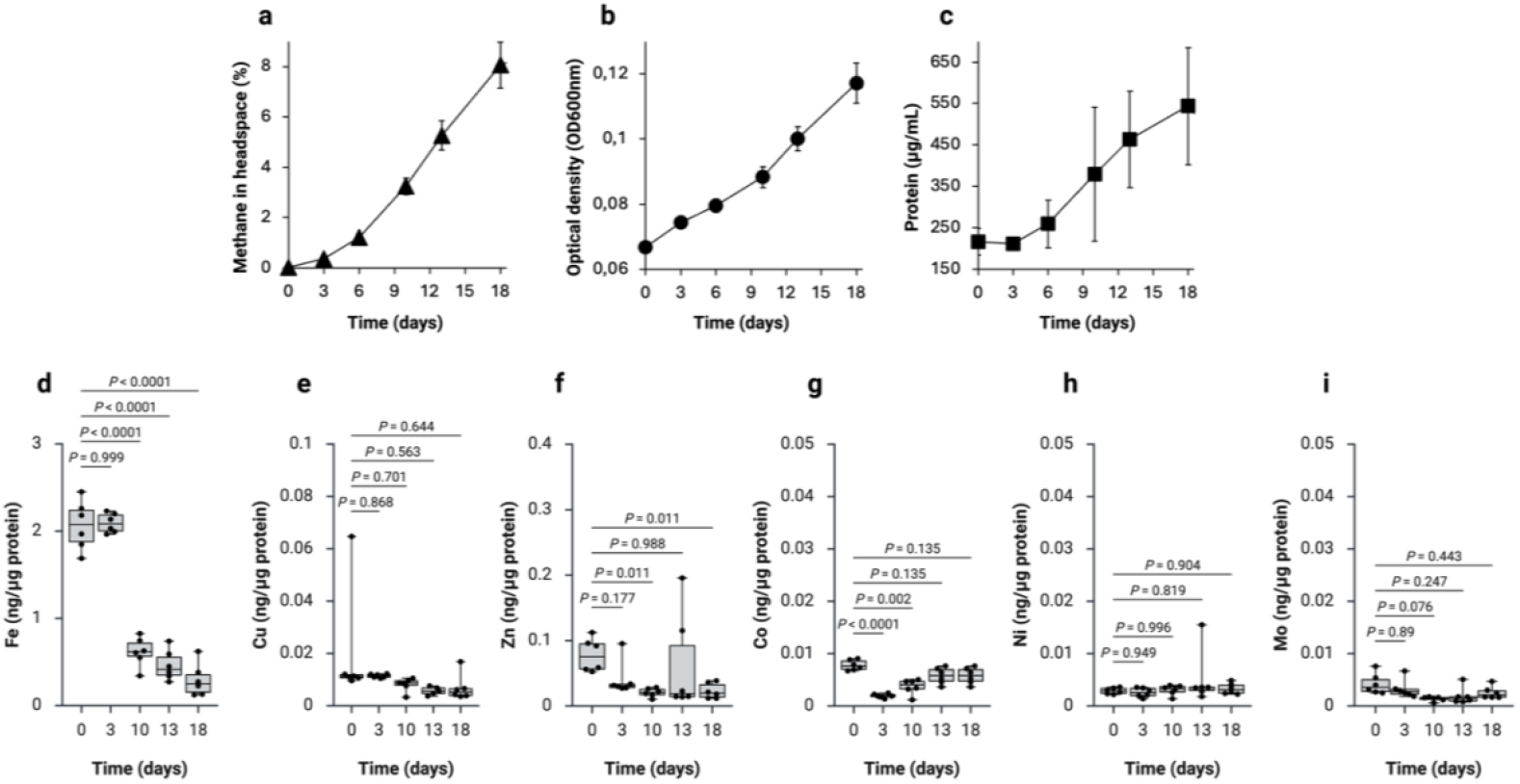
Growth and metal profiles of *M. barkeri* during cultivation with 50 mM acetate. (a) Methane production in the headspace over 18 days, (b) optical density (OD_600nm_) as a measure of cell growth and (c) protein concentration in *M. barkeri* cultures growth on acetate (n=6). (d-i) Concentrations of Fe, Cu, Zn, Co, Ni, and Mo normalized to protein (ng metal/µg protein) over time (n=6). Statistical significance was determined using Welch’s one way ANOVA comparing the initial timepoint to subsequent timepoints to assess metal accumulation during grown on acetate. Figure compiled in Biorender.

### Cellular localization of metals in acetate-grown *Methanosarcina barkeri*

To interpret the metal enrichment observed during electromethanogenesis, we asked where metals reside within the cellular architecture of *M. barkeri*. Three non-exclusive mechanisms could contribute to elevated metal quotas under cathodic growth: (i) redox-triggered shifts in metal homeostasis and uptake, (ii) increased demands for metal cofactors during cathode driven CO_2_-reductive methanogenesis, and (iii) adsorbption/sequestration of metals within the extracellular capsule. Of these, only (iii) can be directly evaluated with subcellular metal mapping. Accordingly, the mapping data presented below most strongly support capsule-associated sequestration as a contributor to some elevated metal quotas, whereas increased cofactor demand and redox-regulated uptake remain plausible, but were not directly tested here.

Because the conductive carbon felt used for the electrodes interferes with ultrathin sectioning and elemental imaging, cathode-attached biomass could not be analyzed by STEM-EDS or nano-XRF. Both elemenal mapping techniques require (depending on the technique) thin, non-conductive, semi-electron-transparent specimens and do not readily resolve the chemistry of the cell layer from that of the underlying felt-electrode surface. We therefore restricted spatial mapping to acetate-grown aggregates and cell capsule and performed these measurements on a limited number of biological replicates due to instrument constraints. Despite this limitation, independent methods yielded consistent evidence of sequestration of specific metals on the extracellular matrix, supporting robust qualitative trends. Nevertheless, the ICP-MS results (section above) provided a comprehensive quantitative dataset spanning all growth conditions, including cathode-associated and control fractions.

This focus on the extracelluar matrix is motivated by the physical organization of *M. barkeri*. The methanochondroitin capsule surrounding *M. barkeri* provides a large extracellular metal-binding compartment (Fig. 4). Electron microscopy of acetate-grown cells shows that M. barkeri forms multicellular aggregates with cells encased in a thick methanochondroitin capsule of 70 – 400 nm, matching earlier estimates by TEM [57] and atomic-force microscopy [58]. Because samples are fixed and dehydrated prior to imaging, the hydrated capsule *in vivo* is likely thicker. A barrier of this magnitude between the extracellular environment and the cytoplasmic membrane implies that any extracellular redox-active component including metals potentially involved in electron uptake, must reside within, or transverse, the capsule. Comparable extracellular matrices in electroactive microorganisms host redox conduits, such as multiheme c-type cytochromes in Gram-negative bacteria where metal cofactors (hemes) are required [59, 60], or flavinylated proteins in some Gram-positives, which do not require metal cofactors [61], while cable bacteria employ Ni-S metalloproteins, with nickel being essential for the nanowires’ conductivity [62].

**Figure 4.**
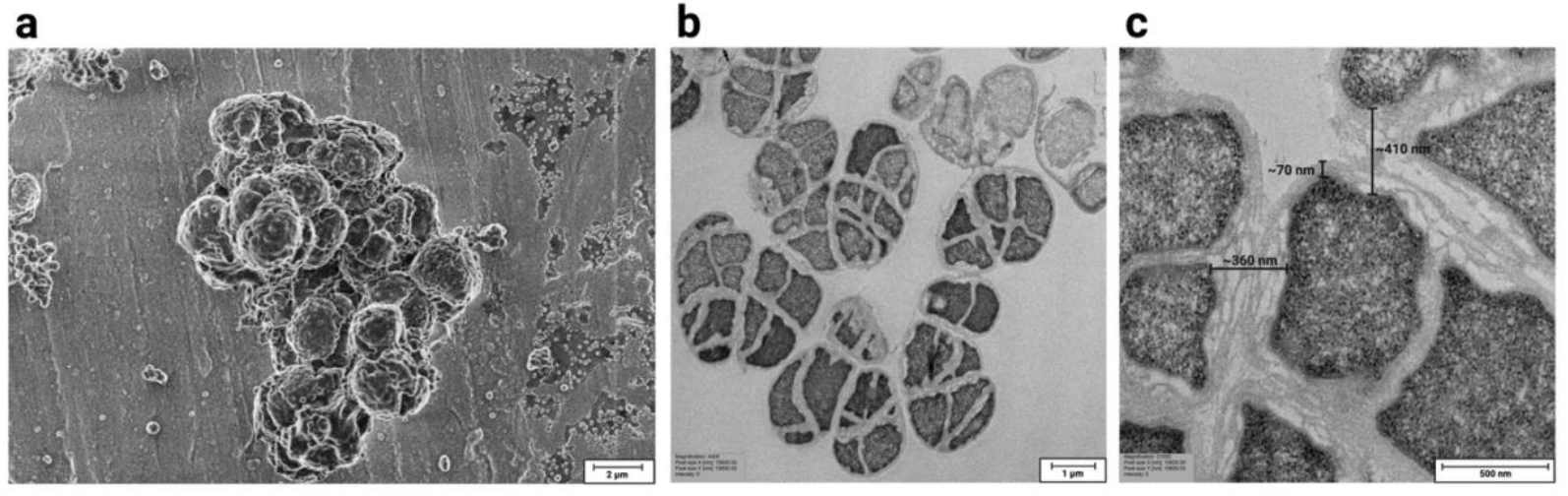
Electron microscopy of acetate-grown *Methanosarcina barkeri* aggregates. (a) Scanning electron micrograph showing a cluster of *M. barkeri* cells encased in methanochondroitin. (b) Transmission electron micrographs of cross sections, illustrating clusters of cells within the methanochondroitin matrix. (c) High magnification transmission electron micrograph highlighting the methanochondroitin coat, with a thickness of ∼70 nm on individual cells and up to ∼400 nm between adjacent cells. Figure compiled in Biorender.

To localize metals within the cellular and capsule compartments of *M. barkeri*, we next used STEM–EDS on ultrathin sections of acetate-grown *M. barkeri* (Fig. 5). This approach mapped Fe primarily intracellularly, while Co and Zn were elevated in the interstitial space between cells, region dominated by methanochondroitin and associated extracellular substances, including G4 nucleic acids [24]. STEM-EDS revealed discrete intracellular hotspots of Fe that co-localized with phosphorus and carbon (Fig. 5a-c), reminiscent of polyphosphates and cyclic 2,3-diphosphoglycerate inclusions previously described in *Methanosarcina frisia* [63]. In contrast, Co, Zn, Mo signals were more diffuse across the section (Fig. 5e-g), and EDS spectra collected from the space between cells (interstitial space likely consisting of methanochondroitin), showed more prominent Co and Zn peaks relative to whole cell regions (Fig. 5i-j), indicating preferential sequestration of these metals in the extracellular matrix.

**Figure 5.**
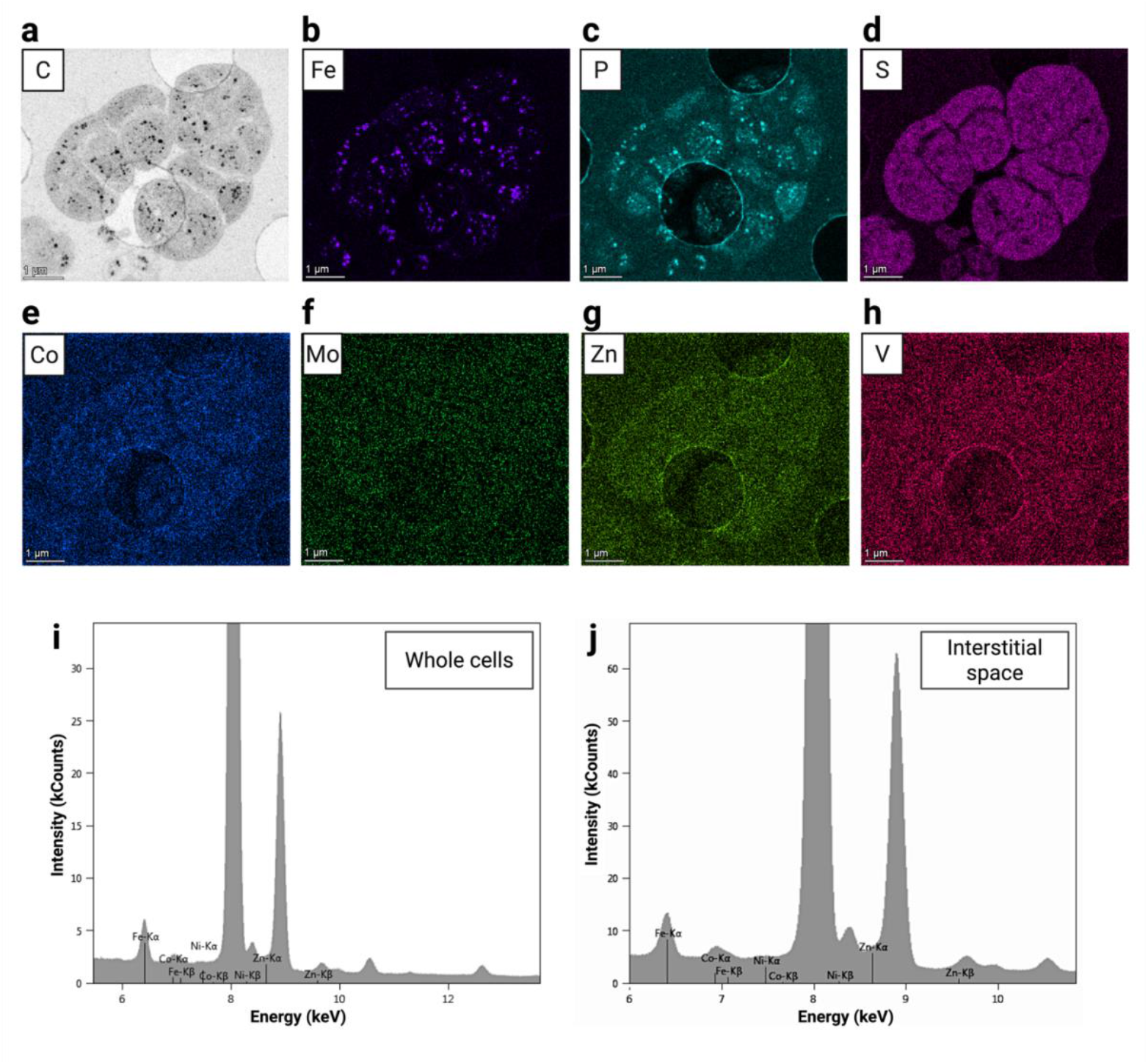
Subcellular localization of metals in acetate-grown *Methanosarcina barkeri* by STEM–EDS. **a)** Scanning-transmission electron micrograph of ultrathin cell sections showing C storage. (b-h) False-colour elemental maps for Fe (violet), P (cyan), S (magenta), Co (blue), Mo (green), Zn (yellow green) and V (red). **i)** Representative EDS spectrum from a whole-cell region highlighting Kα peaks for Fe (6.4 keV), Co (6.9 keV), Ni (7.5 keV) and Zn (8.6 keV). **j)** EDS spectrum from the interstitial space between cells in the aggregate, showing prominent Zn and Co peaks relative to cellular regions, indicatingenrichment of these metals on the extracellular capsule. Scale bars, 1 µm. Figure compiled in Biorender.

To map metals on an extracellular capsule fraction, we applied nanobeam X-ray fluorescence (nano-XRF) [64] on a cell wall fragment from acetate-grown cells (Fig. 6). In this approach, samples are raster-scanned with a ∼50 nm X-ray beam and element-specific fluorescence is collected to generate high-resolution maps. The nano-XRF maps confirmed Co and Zn signals within the capsule material (Fig. 6e-f), consistent with the STEM-EDS observation that these metals are sequestered by the methanochondroitin matrix. In addition, nano-XRF detected Fe associated with the isolated methanochondroitin material. STEM–EDS localized Fe most strongly to intracellular inclusions, likely reflecting the dominance of Fe-rich intracellular hotspots in thin sections compared with the isolated extracellular material analyzed by nano-XRF. Together, these complementary datasets show that the methanochondroitin capsule can sequester Co and Zn (and, to a lesser extent, Fe) providing a mechanistic basis for the elevated metal quota of these metals observed under electromethanogenesis (Fig. 1; Table S1).

**Figure 6.**
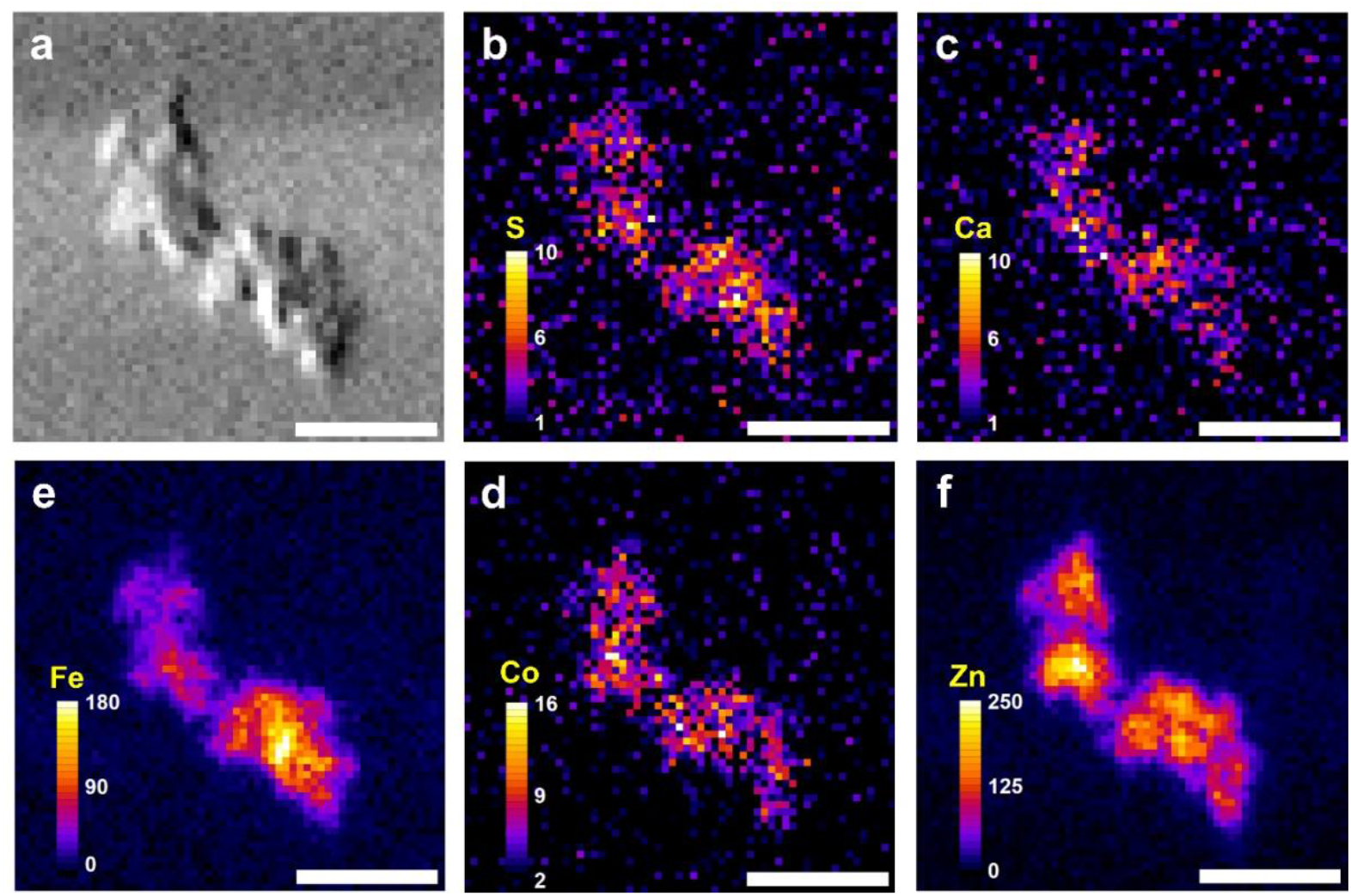
Nano-XRF mapping of a cell surface fragment from *Methanosarcina barkeri*. **(a)** Phase gradient image of a separated methanochondroitin section. (b-f) Elemental distribution maps of Kα emissions for S, Ca, Fe, Co and Zn. Color scales indicate X-ray fluorescence intensity (counts per pixel). Scale bar 1 µm.

### Implications of spatial mapping on biocathode-metal enrichment

Extracellular polymeric substances (EPS), can chelate divalent metals and thereby influence metal homeostasis, biofilm integrity, and extracellular electron transfer [65, 66]. In *M. barkeri*, the spatial mapping of metals, showed Co and Zn (and, to a lesser extent, Fe) associated with the capsule which is a very large methanochondroitin layer, suggesting that this capsule is a substantial extracellular reservoir for these metals. This provides an explanation for the elevated metal quotas (for all three metals) measured by ICP-MS during electromethanogenesis. It is possible that the enrichment of bulk “cell-associated” can arise not only from increased metal uptake inside the cells for cofactor biosynthesis and metabolism, but also from sequestration within the extracellular matrix.

Surface-associated metals in methanogens have been rarely quantified but early work reported higher Zn, Cu, Ni and Fe content on *Methanothrix* sheets relative to *Methanospirillum* [67], consistent with cell matrices acting as a metal reservoir. Complementing these observations, bulk metal quotas have also been rarely quantified. For *M. barkeri*, previous studies have looked primarily at the effect of Ni presence/absence and demonstrated that *M. barkeri* exhibits a high cellular Ni content when it is available and strong Ni-dependent effects on methanogenic performance, linking metal-availability and methanogen physiology[36]. Here we show that during electromethanogenesis metals involved in energy metabolism (Fe, Co, Mo) were significantly enriched in bulk on cells performing electromethanogenesis, and so was Zn which is likely involved in cell envelope quality control/stress responses, and could be consistent with capsule remodelling (attachment/disperssion) under cathodic growth.

Cu represents a significant exception. Although Cu is not involved in the energy metabolism of this methanogen, ICP-MS identified it as the most abundant metal associated with poised biocathode biomass, raising the possibility that Cu participates in electron-transfer at the cell-electrode interface during electromethanogenesis. However, Cu could not be confidently assigned to cellular versus capsule compartments by STEM–EDS or nano-XRF due to method-specific constraints (e.g., Cu-containing grids and limited sensitivity for Cu under our mapping conditions). We therefore interpret Cu enrichment using genomic context (Cu-enzymes found in the genome) and established Cu coordination chemistry. Specific Cu enrichment by cells on poised cathodes is likely due to sequestration onto the cell capsule and possibly by the G4 nucleic acids embedded in it [24]. Because Cu has been shown to stabilize G4-structures, and induce specific folding, and conformational states [49, 50], Cu-could ready G4-structures for electron uptake in *M. barkeri* [24]. The importance of this mechanism in other biological systems remains to be determined.

It is intriguing that cathode-grown *M. barkeri* acts as a strong metal bioaccumulator, concentrating multiple transition metals onto biomass. This raises the possibility that, under specific growth conditions (e.g., direct interspecies electron transfer/DIET[68]), *M. barkeri* cells could act as a local metal reservoirs, buffering trace metal availability for nearby syntrophic partners. More broadly, selective metal enrichment at the cell-material interface could influence how *Methanosarcina* interacts with redox-active materials in its surroundings (e.g., iron minerals, conductive particles, steel surfaces, electrodes)[69, 70]. Importantly, this bioaccumulation phenotype provides a starting point for engineering the electrode-cell microenvironment in bioelectrochemical systems that use methanogens a to upgrade CO_2_ from biogas to methane.

## Conclusion

Our integrated metallomic, imaging and electrochemical analyses show that *M. barkeri*’s interaction with a poised cathode triggers a multifaceted remodeling of its metal economy. During electromethanogenesis, cathode-associated biomass selectively enriched transition metals central for energy metabolism (Fe, Ni, Co and Mo), cell surface turnover (Zn) and coordination of extracellular matrix constituents for electron transfer (e.g., Cu coordinated with G4-RNA) across the thick cell capsule of *M. barkeri*. Together these results suggest that this cytochrome-free methanogen couples intercellular metal cofactor demand with capsule-associated metal sequestration to support electron uptake from a cathode. Beyond establishing metal quotas as a powerful proxy for interrogating archaeal EET strategies our work motivates targeted tests for metal-dependent surface conduits (including metal-G4 nucleic acid interactions), and points to actionable levers (metal availability, capsule-associated sequestration) for optimizing biocathode performance in bioelectrochemical powere to gas technologies.

## Acknowledgment

This article is a contribution to a Novo Nordisk Ascending Investigator grant NNF21OC0067353 awarded to AER. Damien Faivre (D.F.) acknowledges the ERA Chair scheme of the Widening program of the EU for funding of the BioMagnetLink project (grant agreement ID: 101187789). We would like to thank Dr. Satoshi Kawaichi and Dr. Rhitu Kotoky for their advice with electrochemistry experiements, and hydrogen determination tests. We would like to acknowledge the staff at the I14 hard X-ray nanoprobe of Diamond Light Source for technical support. We acknowledge the awarded synchrotron beamtimes under the long term proposal MG28688. We thank the Laboratoire de Maîtrise de la Contamination, de la Chimie des Caloporteurs et du Tritium (LMCT) at CEA Cadarache (DES/IRESNE) for scanning electron microscopy access.

